# Angiotensin II receptor inhibition ameliorates liver fibrosis and enhances hepatocellular carcinoma infiltration by effector T cells

**DOI:** 10.1101/2023.03.05.531188

**Authors:** Li Gu, Yahui Zhu, Maiya Lee, Albert Nguyen, Nicolas T Ryujin, Jian Huang, Shadi Chamseddine, Lianchun Xiao, Yehia I. Mohamed, Ahmed O. Kaseb, Michael Karin, Shabnam Shalapour

## Abstract

Although viral hepatocellular carcinoma (HCC) is declining, non-viral HCC, which often is the end-stage of non-alcoholic or alcoholic steatohepatitis (NASH, ASH), is on an upward trajectory. Immune checkpoint inhibitors (ICI) that block the T cell inhibitory receptor PD-1 were approved for treatment of all HCC types. However, only a small portion of HCC patients show a robust and sustained response to PD-1 blockade, calling for improved understanding of factors that negatively impact response rate and duration and the discovery of new adjuvant treatments that enhance ICI responsiveness. Using a mouse model of NASH-driven HCC, we identified peritumoral fibrosis as a potential obstacle to T cell mediated tumor regression and postulated that anti-fibrotic medications may increase ICI responsiveness. We now show that the angiotensin II receptor inhibitor losartan, a commonly prescribed and safe antihypertensive drug, reduced liver and peritumoral fibrosis and substantially enhanced anti-PD-1 induced tumor regression. Although losartan did not potentiate T cell reinvigoration, it substantially enhanced HCC infiltration by effector CD8^+^ T cells compared to PD-1 blockade alone. The beneficial effects of losartan correlated with inhibition of TGF-β receptor signaling, collagen deposition and depletion of immunosuppressive fibroblasts.

**Significance:** Immune checkpoint inhibitors are used in HCC treatment but overall response rates for single agent PD-1/PD-L1 blockers have remained stubbornly low. Using a mouse model of NASH-driven HCC, we show that co-treatment with the safe and inexpensive angiotensin II receptor inhibitor losartan substantially enhanced anti-PD-1 triggered HCC regression. Although losartan did not influence the reinvigoration of exhausted CD8^+^ T cells it considerably enhanced their intratumoral invasion, which we postulated to be compromised by peritumoral fibrosis. Indeed, the beneficial effect of losartan correlated with inhibition of TGF-β signaling and collagen deposition, and depletion of immunosuppressive fibroblasts. Losartan should be evaluated for its adjuvant activity in HCC patients undergoing PD-1/PD-L1 blocking therapy.

## Introduction

Hepatocellular carcinoma (HCC), one of the most common cancer types worldwide (1), is the end-result of chronic liver injury and inflammation, often occurring in the context of hepatocyte cell death and liver fibrosis (2, 3). Whereas early and locoregional HCC are effectively treated by surgical resection, radiofrequency ablation or chemoembolization, treatment options for advanced HCC are limited by the compromised liver function accompanying the disease (1). The only approved targeted HCC therapies are pan-kinase inhibitors, such as sorafenib, which extend patient survival by several months, leaving the 5-year survival rates at 30% for patients with localized disease and an abysmal 2.5% for patients with advanced metastatic disease (4, 5). A considerable advance in HCC treatment was the finding that immune checkpoint inhibitors (ICI) targeting PD-1 or its ligand PD-L1 [hereafter PD-(L)1], whose association causes T cell exhaustion (6), can curtail HCC growth or induce tumor regression with objective response rates of 15-20% (7-11). These rather modest response rates were recently improved by combining PD-(L)1 blockade with VEGF receptor (VEGFR) inhibition or CTLA4 checkpoint blockade (11). Despite these improvements, response rates remain lower than 30% and suggested to be particularly low in non-alcoholic steatohepatitis (NASH)-related HCC (12-14). Although hepatosteatosis was postulated to account for the adverse effect of NASH on ICI responsiveness (12-14), it should be recognized that hepatosteatosis usually declines or disappears (“burnout NASH”) in advanced NASH, which is characterized by extensive fibrosis preceding HCC development (15). Moreover, hepatosteatosis is not required for induction of liver damage and fibrosis in NASH afflicted mice (16). Based on our studies of NASH-related HCC in the *MUP-uPA* mouse model (17-19), we reasoned that liver fibrosis is more likely to account for the adverse effect of NASH on ICI outcome than hepatosteatosis. While NASH is driven by metabolic inflammation, it is also accompanied by marked changes in the hepatic immune system, including the accumulation of immunosuppressive IgA-expressing plasma cells, which dismantle immunosurveillance by HCC-directed T cells (19, 20). The immunosuppressive activity of IgA^+^ plasma cells is IL-10 and PD-L1 dependent and either PD-L1 or IL-10 blockade, or ablation, restore anti-HCC immunity to high fat diet (HFD) fed *MUP-uPA* mice (19), which develop NASH and robustly progress to HCC (17, 18). While most NASH induced liver tumors in *MUP-uPA* mice were effectively eliminated by PD-L1 blockade, tumors with extensive peritumoral fibrosis were treatment refractory (19).

Liver fibrosis or excessive collagen fiber deposition is triggered by chronic liver injury, which induces production of TGF-β and other pro-fibrogenic cytokines by activated immune cells, mainly macrophages (21, 22). TGF-β activates collagen-producing hepatic stellate cells (HSC) that express α smooth muscle actin (αSMA) and glial fibrillary acidic protein (GFAP) (23) and gives rise to cancer associated fibroblasts (CAF) during HCC emergence (24, 25). TGF-β activates HSC by binding to its type II receptor (TGFBR2) which heterodimerizes with the type I receptor (TGFBR1), triggering activation and nuclear translocation of SMAD2, 3 and 4 transcription factors (26). TGFBR signaling is potentiated by angiotensin II (Ang II) acting via its type 1 receptor (AngIIR1) (27, 28). Although no TGFBR inhibitors or other targeted therapeutics were approved for the treatment of liver fibrosis (29), the commonly used AngIIR1 inhibitor and antihypertensive drug losartan can reduce liver fibrosis in humans (30, 31) and rodents (32, 33). Studies by Rakesh Jain’s group have further shown that losartan enhances cancer drug delivery (34) and can downregulate immunosuppression-associated genes in ovarian and pancreatic cancers when combined with chemo-or radio-therapy (35, 36). Inspired by these findings, we investigated if losartan improves ICI-induced HCC regression and if so, whether this correlates with its antifibrogenic activity. We now show that losartan potentiates the therapeutic response to a suboptimal PD-1 antagonistic antibody in the *MUP-uPA* model and that this effect correlates with improved intra-tumoral invasion by re-invigorated CD8^+^ cytotoxic T cells (CTLs), diminished collagen type I (Col I) production and downregulated TGF-β signaling.

## Results

### Losartan potentiates anti-PD-1 induced HCC regression

To determine whether losartan can potentiate anti-PD-1 induced HCC regression, we placed 6-weeks old *MUP-uPA* mice on HFD for 6 months to induce NASH and HCC. HCC bearing mice, still maintained on HFD, were then divided into 4 groups (*SI Appendix*, Fig. S1*A)* that were given control (ctrl) IgG, anti-PD-1, losartan + ctrl IgG, or losartan + anti-PD-1 for 8 weeks. Body weight gain was identical across treatment groups and no organ injury was observed (*SI Appendix*, Fig S1 *B* and *C*). Notably, the combination of losartan with anti-PD-1 resulted in lower liver/body weight ratio, tumor multiplicity and tumor volume compared to ctrl IgG, anti-PD-1 alone or ctrl IgG plus losartan (Fig. 1 *A-D*). Importantly, losartan addition augmented anti-PD-1 induced tumor regression. However, anti-PD-1 single treatment also caused a moderate decrease in hepatosteatosis, liver triglyceride (TG) accumulation and serum TG amounts, effects that were slightly affected by losartan addition (*SI Appendix*, Fig. S1 *D-F*). Losartan without or with anti-PD-1 reduced liver damage marked by the presence of liver enzymes in the circulation (*SI Appendix*, Fig. S1*G*).

**Fig. 1.**
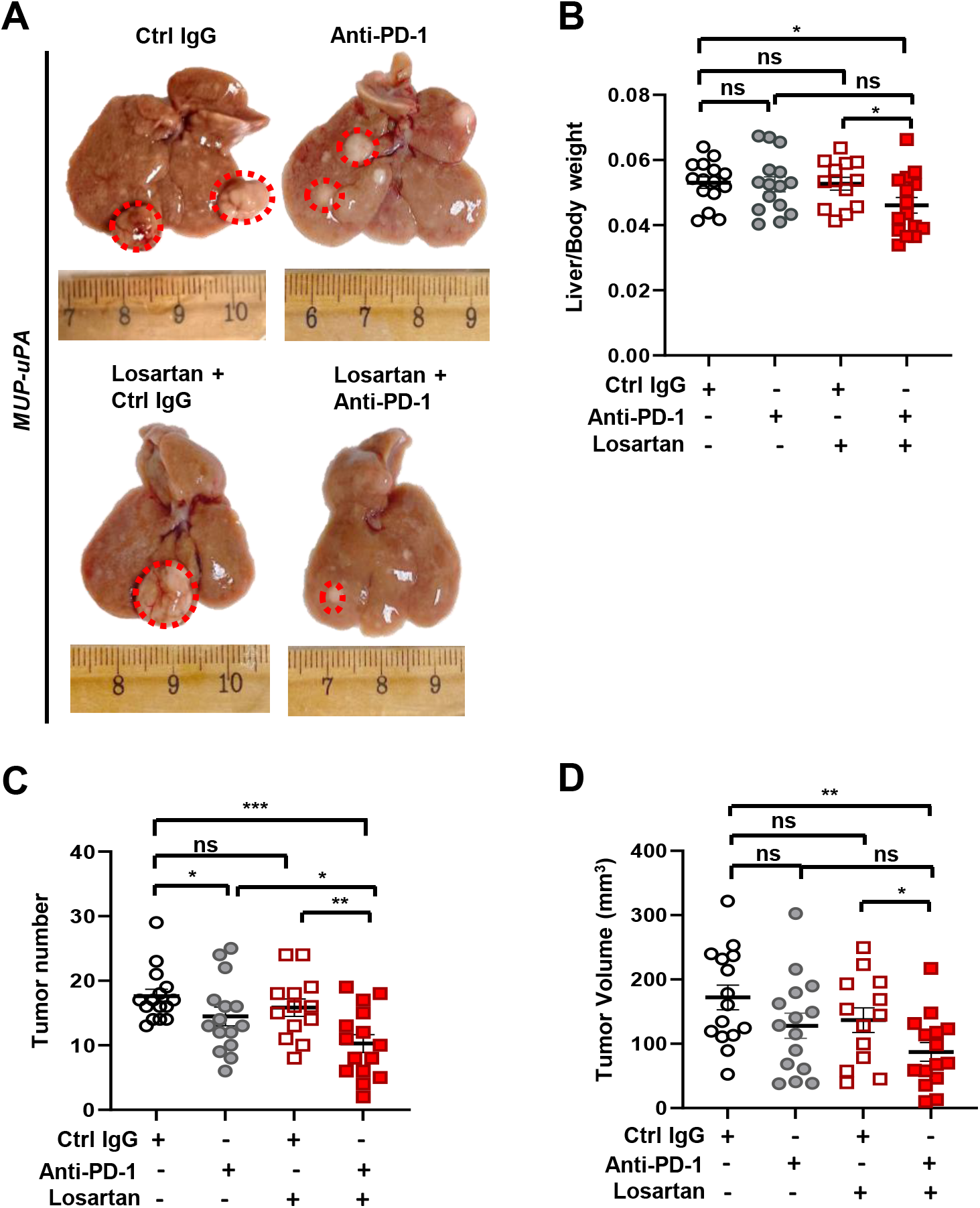
Losartan potentiates anti-PD-1 induced regression of NASH-driven HCC. (*A*) Gross liver morphology in HFD-fed *MUP-uPA* mice that were treated with ctrl IgG, anti-PD-1, losartan + ctrl IgG, and losartan + anti-PD-1 (n=13-15). (*B*) Liver/Body weight ratio in above mice at the end of the treatments. (*C* and *D*) Tumor multiplicity (C) and volume (D) at the end of treatments. Data are presented as mean ± SEM. *P < 0.05, **P < 0.01, ***P < 0.001 (Unpaired two-tailed t test and Mann-Whitney test were used to determine significance).

### Losartan enhances HCC infiltration with anti-PD-1 induced T effector cells

To identify how losartan enhanced HCC regression triggered by anti-PD-1 treatment, immune cell numbers and effector functions were analyzed by flow cytometry (FC). Both CD8^+^ T cell number and fraction were increased in the livers of anti-PD-1 treated mice, with the majority of CD8^+^ T cells secreting higher amounts of TNF and IFNγ (Fig. 2*A, SI Appendix*, Fig. S2 *A*-*C*). Conversely the number of CD8^+^ T cells that express the inhibitory collagen receptor LAIR1 (37) declined after anti-PD-1 treatment (*SI Appendix*, Fig. S2*D*). Losartan, however, had no effect on total CD8^+^ T cell number, effector function or LAIR1 expression (*SI Appendix*, Fig. S2 *A*-*D*). Nonetheless, the combination of losartan with anti-PD-1 increased the number and percentage of CD8^+^ T cells associated with lower tumor volume and number in comparison to anti-PD-1 alone (Fig. 2 *A-C*). Anti-PD-1 without or with losartan enhanced the expression of mRNAs for Ccl8, Ccl5, Ccl19, Ccl2, Cxcl9 and Cxcl10, chemokines that promote T cell recruitment and activation (*SI Appendix*, Fig. S2 *E*-*G*). Losartan co-treatment trended to further increase chemokine expression, but the effect was not statistically significant. FC and immunochemistry (IHC) revealed that while losartan in combination with anti-PD-1 did not exert a significant effect on anti-PD-1 induced CD8^+^ T cell re-invigoration or CD8^+^ T cell activation markers, it increased tumor-infiltration by CD8^+^CD3^+^ T cells, CD3^+^CD8^-^ helper T cells (CD4^+^), CD3^-^CD8^+^ plasmacytoid dendritic (pDC) cells and CD45^+^ immune cells compared to anti-PD-1 alone (Fig. 2 *D-G* and *SI Appendix*, Fig. S2 *H-J*). These results show that the main effect of losartan on anti-HCC immunity was to increase tumor infiltration with re-invigorated T cells and pDC. Losartan co-treatment, however, did not enhance the anti-PD-1 induced expression of MHC-I related genes, which present antigens to CD8^+^ T cells (38), such as *Nlrc5, Psm9*, and *Tap1*, neither did it affect *Il1b* or *Cd274* mRNA expression (*SI Appendix*, Fig. S2 *K-O*).

**Fig. 2.**
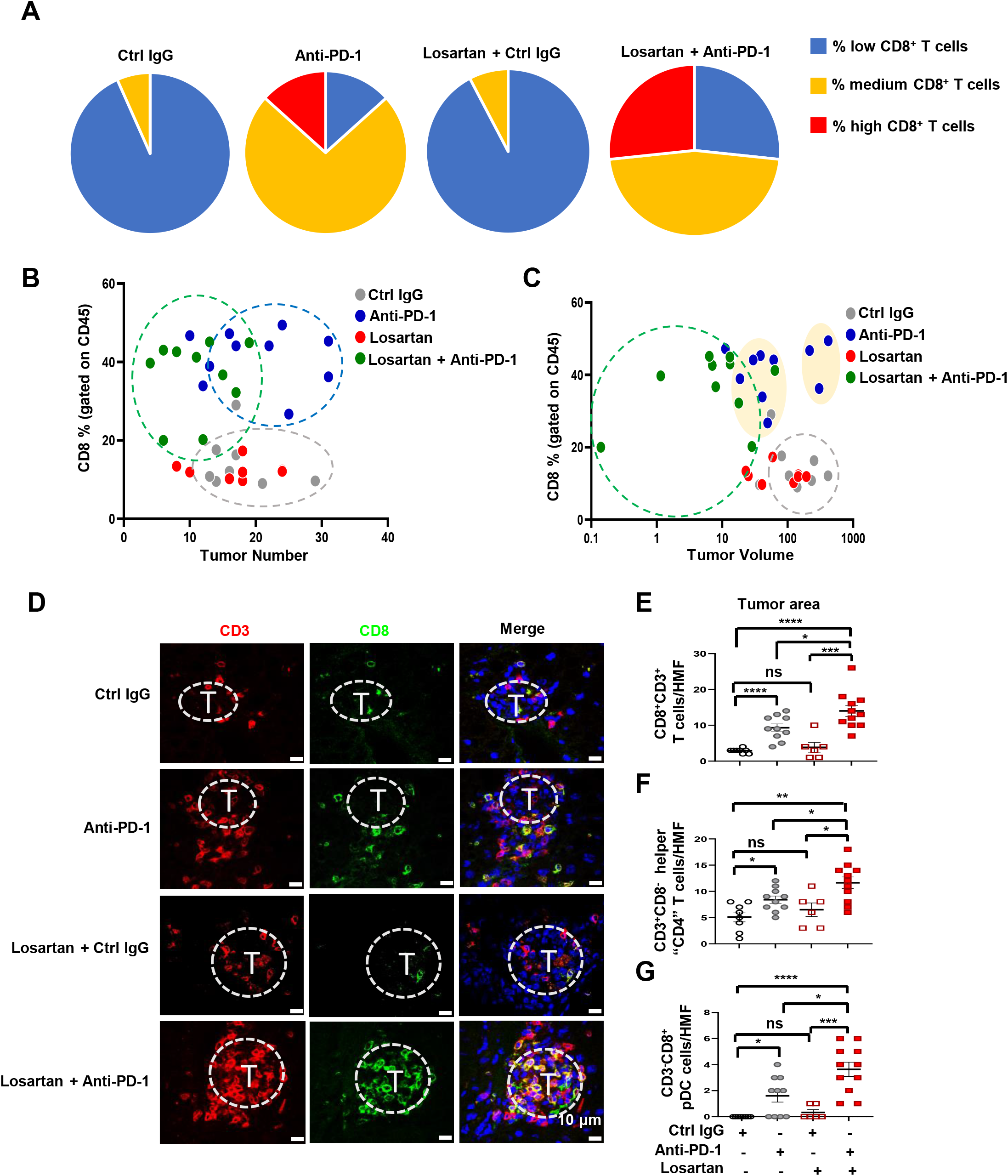
Losartan stimulates intratumoral infiltration by re-invigorated Teff cells. (*A*) *MUP-uPA* mice from each group in Fig. 1A were classified into 3 categories based on the number of CD8^+^ T cells in their livers (low<1million; medium 1 to 4.6 million; high>4.6 million). The percentages of mice in each category are indicated (n=13-15). (*B* and *C*) Correlation between liver CD8^+^ T cell frequency and treatment outcome determined as either tumor multiplicity (B) or tumor volume (C). (*D-G*) Frozen liver sections (n=6-8 per mouse) were stained for CD3 and CD8, and counterstained with DAPI. T-tumor. Scale bars, 10 µm (D). Quantification of CD8^+^CD3^+^ T cells (E), CD3^+^CD8^-^ CD4^+^ helper T (F) cells and CD3^-^CD8^+^ pDC (G) into tumors is based on cell quantitation per high magnification field (HMF) in tumor and non-tumor areas by Image J analysis of 10 fields of per section. Data are presented as mean ± SEM. *P < 0.05, **P < 0.01, ***P < 0.001, ****P < 0.0001 (Unpaired two-tailed t test and Mann-Whitney test).

### Losartan inhibits liver fibrosis

We next explored likely mechanisms through which losartan stimulates HCC infiltration by T cells. Losartan alone or together with anti-PD-1 largely reduced liver fibrosis, assessed by Sirius Red staining, which was slightly increased by anti-PD-1 alone (Fig. 3 *A and B*). The major extracellular matrix (ECM) protein collagen type I α1 chain (Col1a1) and the HSC marker α-SMA also declined after losartan alone or losartan + anti-PD-1 (Fig. 3 *A and C-G*). Losartan, however, had a modest effect on fibroblast-specific protein 1 (FSP1) expressing fibroblasts (Fig. 3 *E* and *H*). Unlike α-SMA^+^ HSC, FSP1^+^ fibroblasts support the response to immunotherapy by producing chemokines (39). Of note, losartan co-treatment increased the numbers of lymphoid dense areas (tertiary lymphoid follicle-like structures, TLFS), which contained FSP1^+^ fibroblasts next to CD8^+^ T cells and B220^+^ B cells (*SI Appendix*, Fig. S3 *A*-*C*), and previously shown to predict better prognosis (40). COX2^+^α-SMA^+^ fibroblasts and COX2^+^FSP1^+^ fibroblasts, which have immunosuppressive properties (41), were lower after losartan treatment (*SI Appendix*, Fig. S3 *D-G*). Altogether, losartan treatment reduced liver fibrosis, inhibited Col1a1 deposition and blunted the generation of immunosuppressive CAF.

**Fig. 3.**
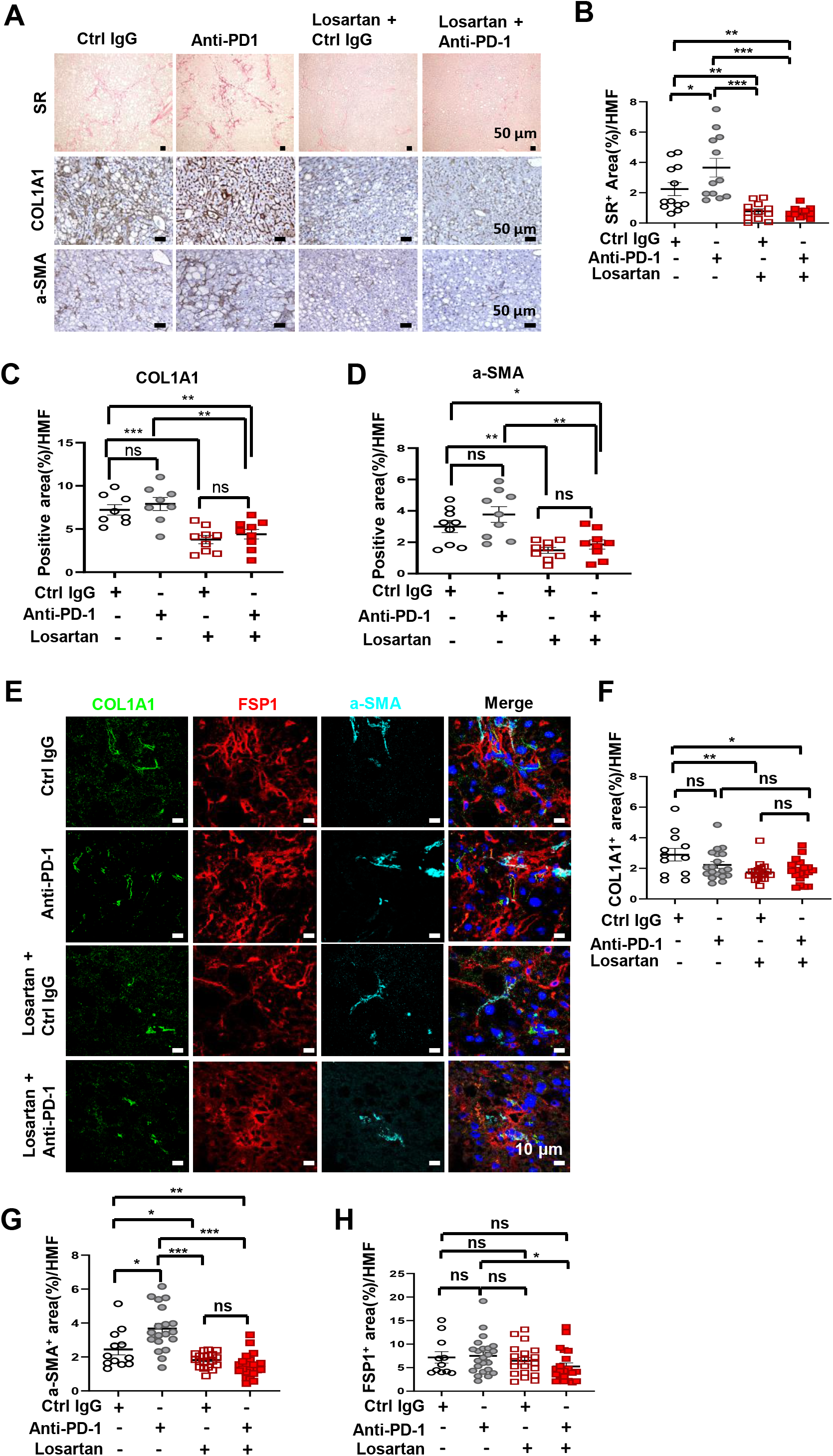
Losartan inhibits NASH-related liver fibrosis. (*A*) FFPE liver sections were stained with Sirius Red (SR) (top), collagen I alpha 1 chain (COL1A1) (middle), and α-SMA (bottom) antibodies (n=8). Scale bars, 50 µm. (*B-D*) SR (B), COL1A1 (C), and α-SMA (D) staining intensities per HMF were determined by Image J quantitation of 10 fields per section. (*E*) Frozen liver sections were stained for COL1A1, FSP1, and α-SMA and examined by fluorescence microscopy. Scale bars, 10 µm. Experiments were repeated at least three times. (*F-H*) Quantification of COL1A1^+^ (F), α-SMA^+^ (G), and FSP1^+^ (H) areas per HMF. Data are presented as mean ± SEM. *P < 0.05, **P < 0.01, ***P < 0.001 (Unpaired two-tailed t test and Mann-Whitney test).

### Losartan inhibits TGF-β signaling

Next, we examined the effect of losartan on TGF-β signaling. IHC showed that losartan inhibited ERK1/2 phosphorylation in hepatocytes and stellate cells, as well as TGF-β1 expression (Fig. 4 *A* and *B*). Immunoblotting (IB) confirmed the decrease in ERK1/2 phosphorylation and showed that losartan also inhibited SMAD2 and 3 phosphorylation (Fig *4C*). Q-RT-PCR showed that losartan also blunted *Tgfbr1, Tgfbr2, Vegf, Pdgfb, Pdgfrα, Pdgfrβ, Ctnnb1* (β-Catenin) and *Fgf2* mRNA expression (*SI Appendix*, Fig. S4 *A* and *B*). Expression of the TGF-β target connective tissue growth factor (CTGF) also decreased after losartan without or with anti-PD-1 (*SI Appendix*, Fig. S4 *A, C* and *D*). In agreement with a recent publication, anti-PD-1 treatment increased IL-6 expression (Fig. 4*D* and *SI Appendix*, Fig. S4*E*), which supports ICI resistance (42). Losartan co-treatment, however, reversed this effect and reduced the number of IL6^+^α-SMA^+^ fibroblasts (Fig. 4*D* and *SI Appendix*, Fig. S4*E*). Consistent with the ability of PD-1 blockade to improve senescence surveillance (43), anti-PD-1 treatment reduced p21 and p16 expression, an effect that was modestly enhanced by losartan co-treatment (Fig. 4 *E*-*G*). Anti-PD-1 increased expression of hexosamine pathway (HBP) genes (*SI Appendix*, Fig. S4 *F-J*), including glutamine-fructose-6-phosphate transaminase 1 (*Gfpt1*), O-linked N-acetylglucosamine (*GlcNAc*) transferase (*Ogt*), phosphoglucomutase 3 (*Pgm3*), UDP-N-acetylglucosamine pyrophosphorylase 1 (*Uap1*) and the EGFR ligand amphiregulin (*Areg*), which promotes ECM hyaluronan synthesis (44). The addition of losartan, however, reduced the expression of these genes (*SI Appendix*, Fig. S4 *F-J*). Collectively, these results demonstrate that losartan has a profound effect on the ECM and the tumor stroma, effects that are consistent with the inhibition of TGF-β signaling and likely to contribute to enhancement of tumor invasion by CD8^+^ Teff cells.

**Fig. 4.**
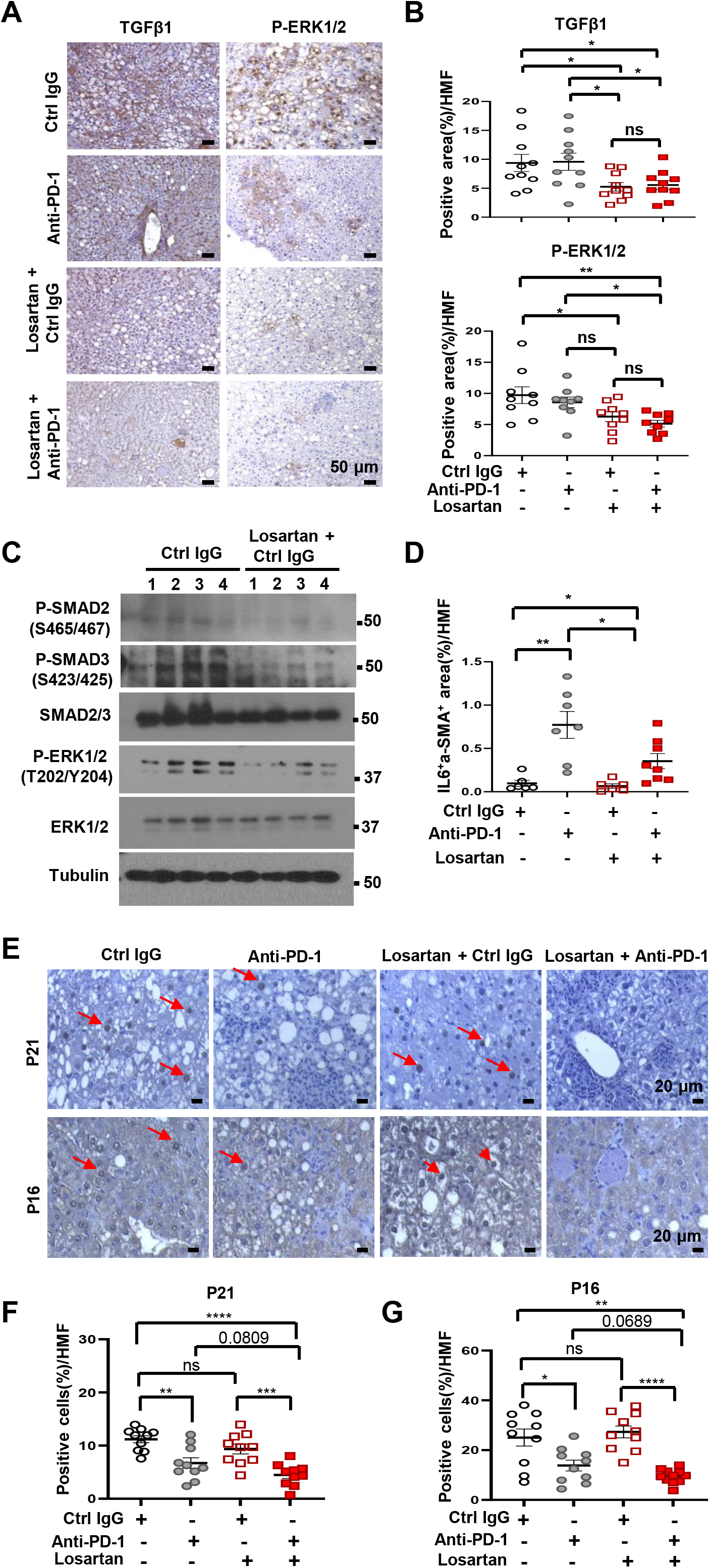
Losartan promotes stromal remodeling by inhibiting TGF-β expression and signaling. (*A*) IHC staining of TGF-β1 (left) and phosphorylated (P)-ERK1/2 (right) in FFPE liver sections from *MUP-uPA* mice in the different treatment groups (n=8). Scale bars, 50 µm. (*B*) TGF-β1 (upper) and P-ERK1/2 (bottom) staining intensity per HMF determined by Image J. (*C*) Immunoblot analysis of the indicated proteins in liver lysates of mice from the indicated treatment groups (n=4). (*D*) Quantification IL6^+^α-SMA^+^ area per HMF by Image J analysis of the data in Fig. S4E. (*E-G*) IHC staining of p21 and p16 in FFPE liver sections from the indicated treatment groups (n=8) (E). Scale bars, 20 µm. p21 (F) and p16 (G) positive cells per HMF determined by Image J. Data are presented as mean ± SEM. *P < 0.05, **P < 0.01, ***P < 0.001, ****P < 0.0001 (Unpaired two-tailed t test and Mann-Whitney test).

## Discussion

Our results show that losartan, a safe, inexpensive, and widely used AngIIR1 antagonist, significantly potentiates HCC regression in response to PD-(L)1 blockade. Of note, losartan had no effect on the activation state of hepatic T cells and their expression of the inhibitory collagen receptor LAIR1, on its own or together with anti-PD-1. The only obvious effect of losartan co-treatment on anti-HCC immunity was the enhancement of HCC infiltration by CD8^+^ T cells that were reinvigorated by PD-1 blockade, as well as by pDC. Without added losartan, anti-PD-1 treatment resulted in the expected increase in Teff cells, but the reinvigorated CD8^+^ T cells mainly remained at the tumor margin with very few of them detected within the tumor proper. Consistent with previous publications (32, 33), losartan treatment alone or in combination with anti-PD-1 ameliorated liver and peritumoral fibrosis, an effect that was likely due to inhibition of TGFBR1, TGFBR2 and CTGF expression and SMAD and ERK phosphorylation, as well as diminished Col I production due to inhibition of HSC activation, all of which reflect inhibition of TGF-β signaling. Indeed, the effects of losartan closely resemble the anti-PD-L1 potentiating effect of TGF-β1 blockade in a mouse model of colorectal cancer (45). These results are consistent with our previous finding that HCCs that were surrounded by more extensive peri-tumoral fibrosis were refractory to PD-L1 blockade compared to HCC nodules lacking a fibrotic envelope (19). Although it was already reported that losartan inhibits Col I production and improves blood vessel perfusion, leading to improved drug delivery and enhanced chemotherapy and radiation effectiveness (34, 46, 47), this is the first time that losartan was shown to increase ICI responsiveness. Our results were obtained in a mouse model of NASH-driven HCC that shares many features with the equivalent human disease (17). Nonetheless, it is important to conduct retrograde analysis of human clinical data and determine if HCC patients who received losartan or other AngIIR1 inhibitors exhibit an improved ORR when subjected to PD-(L)1 blockade. Moreover, it needs to be tested whether losartan can potentiate the response to PD-(L)1 blockade in other cancers associated with peritumoral fibrosis and desmoplasia, such as pancreatic cancer and intrahepatic cholangiocarcinoma, which so far have been refractory to ICI. As losartan only slightly affect hepatosteatosis, our results suggest that liver fibrosis maybe a more relevant explanation of the modest response of NASH-induced HCC to PD-(L)1 blockers (12-14).

## Materials and Methods

### Animals

*MUP-uPA* mice were previously described and kindly provided by E.P. Sandgren, University of Wisconsin-Madison (48). Mice were maintained in filter-topped cages on autoclaved food and water with a 12 h light (6am-6pm)/ dark (6pm-6am) cycle. To induce NASH and HCC, male mice were placed on HFD (high fat Diet, Bio-Serv S3282) at 6-8 weeks-of-age. After 6 months, the mice were administered control IgG (25AUW, mouse [HEXON-Ad] mAb (TC31.27F11.C2) IgG1 D265A/Kappa) (20 mg/kg, i.p.) or anti-PD-1 (03AHF, mouse modified PD-1 mAb (DX400 D265A LPD2127/LPD2128) mIgG1/Kappa), 2-3 times weekly (20 mg/kg, i.p.) without or with losartan (30 mg/kg in drinking water) (*SI Appendix*, Fig. S1*A*). Mice were monitored weekly, and body weight gain and food consumption were calculated every 2 weeks. After 8 weeks, the mice were euthanized, and tumors and livers analyzed. Tumor volume was calculated as: LxWxH/2 (L, length, W, width, H, height). All experiments were performed according to UCSD Institutional Animal Care and Use Committee and NIH guidelines and regulations. Dr. Karin’s Animal protocol S00218 was approved by the UCSD Institutional Animal Care and Use Committee.

### Flow Cytometry

Single cell suspensions were prepared from livers and spleens. For liver lymphocyte isolation, 0.5 g of tissue was cut into small pieces and incubated in dissociation solution (RPMI medium supplemented with 5% FBS, collagenase type I (200 U/ml), collagenase type IV (200 U/ml), and DNase I (100 μg/ml) for 40 min. at 37°C. Next, the cell suspensions were passed through a 40 μm cell strainer and washed twice. Isolated cells were incubated with labeled antibodies in Cell Staining Buffer (Biolegend). Dead cells were excluded based on staining with Live/Dead Fixable Viability Dye (FVD-eFluor780, eBioscience). For intracellular IgA staining, cells were additionally fixed and permeabilized with BD™ Perm/Wash buffer (BD Biosciences) before being stained with labeled IgA antibody. For intracellular cytokine staining, cells were re-stimulated with cell stimulation cocktail (eBioscience; containing PMA and ionomycin), in the presence of a protein transport inhibitor cocktail (eBioscience; containing brefeldin A and monensin). After 4 hrs. incubation at 37°C, cells were fixed and permeabilized with BD™ Perm/Wash buffer (BD Biosciences). After fixation/permeabilization, cells were stained with labeled antibodies of interest. Cells were analyzed on a Beckman Coulter Cyan ADP flow cytometer. Data were analyzed using FlowJo software (Treestar). Absolute numbers of specific immune cells (e.g., CD8^+^ cells) in spleens were calculated by multiplying the total cell numbers from one spleen by the percentage of the cell type in question among total CD45^+^ immune cells. Absolute immune cell numbers in livers were calculated by multiplying total cell number in one liver fragment by the percentages of the corresponding cell type amongst all total liver cells divided by the weight of the analyzed liver fragment (cell number per gram of liver).

### Histology

Livers were removed, and portions of liver tissue were fixed in 4% paraformaldehyde and embedded in paraffin. Thick sections (5 μm) were stained with hematoxylin and eosin (H&E) (Leica, 3801615, 3801571), Sirius Red (ab246832) and processed for IHC. For frozen block preparations, liver tissue fragments were embedded in Tissue-Tek OCT compound (Sakura Finetek), sectioned and stained with Oil Red O (ORO). Image J was used for image quantification as descried (19). Briefly, for Sirius Red, areas of at least 1 mm^3^ were quantitated with Image J and normalized for vascularization and lipid accumulation using corresponding H&E-stained areas. For ORO analysis, multiple images (3-4) were quantitated and averaged using Image J. IHC was performed as follows: after xylene de-paraffinization and rehydration with ethanol series, antigen retrieval was conducted for 15 min at 100℃ with 0.1% sodium citrate buffer. After quenching of endogenous peroxidases with 3% H_2_O_2_ and blocking with 5% bovine serum albumin (BSA), sections were incubated with indicated antibodies (Table S1) overnight at 4℃ followed by incubation with biotinylated secondary antibodies (1:200) for 30 min and Streptavidin-HRP (1:500) for 30 min. Bound peroxidase was visualized by 1-10 min incubation in 3, 30-diaminobenzidine (DAB) solution (Vector Laboratories, SK-4100). Images were captured on an upright light/fluorescent Image A2 microscope with AxioVision Release 4.5 Software (Zeiss, Germany).

### IB analysis

Livers were homogenized in a Dounce homogenizer (Thomas Scientific, NJ) with 30 strokes in RIPA buffer (50 mM Tris-HCl, pH 7.4, 150 mM NaCl, 1% Triton X-100, 1% sodium deoxycholate, 0.1% SDS, 1 mM EDTA) with complete protease and phosphatase inhibitor cocktail. Lysates were sonicated, centrifuged, and boiled in 4× loading buffer. The samples were separated by SDS-PAGE and transferred to PVDF membranes, blocked in 5% nonfat milk, and incubated with the indicated primary antibodies overnight. Secondary antibodies were added for another 1 hr. and detected with Clarity Western ECL Substrate (Biorad). Immunoreactive bands were exposed in an automatic X-ray film processor. Antibodies are listed in Table S1.

### Immunostaining

Tissues were embedded in Tissue Tek OCT (Sakura Finetek) and snap frozen. Tissue sections were fixed in cold acetone/ methanol for 10 min and washed with PBS. Slides were blocked with PBS/1% normal donkey serum for 30 min. Sections were incubated with primary antibodies overnight at 4℃. After washing with PBS, secondary antibodies and DAPI were added for 1 hr. at room temperature.

Slides were washed with PBS and covered with FluorSave Reagent (EMD Millipore, 345789). Images were captured on a TCS SPE Leica confocal microscope. Results were quantified by counting dots/calculating intensity for each field of view (4-5 areas for each slide) with Image J.

### Metabolic measurements

Liver and serum TG were measured with Triglyceride Colorimetric Assay Kit (Cayman Chemical #10010303) according to manufacturer protocol. Circulating ALT was measured with ALT(GPT) Reagent (Thermo Scientific™, TR71121) according to manufacturer protocol.

### RNA isolation and quantitative real-time PCR (Q-RT-PCR)

Total liver RNA was extracted with RNeasy Plus Mini kit (Qiagen #74134) and cDNA was synthesized with SuperScript™ VILO™ cDNA Synthesis Kit (Thermo Fisher Scientific, 11754050). mRNA amounts were determined on a CFX96 thermal cycler (Biorad). Data are presented as arbitrary units and calculated by the comparative CT method (2Ct^(18s rRNA–gene of interest)^). Primers are listed in Table S2.

### Quantification and statistical analysis

Data are presented as mean ± SEM. Differences between mean values were analyzed by two-tailed student’s t test and Mann-Whitney test with GraphPad Prism software. P value<0.05 was considered as significant (*p < 0.05, **p < 0.01, ***p < 0.001, ****p < 0.0001).

## Data, Materials, and Software Availability

All study data are included in the article and/or SI Appendix.

## Acknowledgements

We thank Cell Signaling Technologies, Santa Cruz Technologies, and Life Technologies for gifts of antibodies/other reagents and the UCSD histology core for assistance. Funding: Research was supported by the Merck and Co. OSTP program and NIH grants to M.K. (R01CA234128) and M.K. and S.S. (U01AA027681). Research was also supported in part by a research grant from Investigator-Initiated Studies Program of Merck Sharp & Dohme LLC. The opinions expressed in this paper are those of the authors and do not necessarily represent those of Merck Sharp & Dohme LLC. Additional support came from the MD Anderson Cancer Center SPORE in Hepatocellular Carcinoma to S.S. (P50 CA217674). M.K. hold the Ben and Wanda Hildyard Chair for Mitochondrial and Metabolic Diseases and is an American Cancer Society Research Professor.

## Author contributions

M.K., and S.S. conceived the project. L.G. and Y.Z. designed the study and performed most of the experiments. M.L., A.N., N.T.R, and J.H. assisted with experiments and data analysis. S.S. provided reagents, advice and helped with FC data interpretation and discussion. M.K., L.G. and S.S. wrote the manuscript, with all authors contributing and providing feedback.

