## supplementary data for "Angiotensin II receptor inhibition ameliorates liver fibrosis and enhances hepatocellular carcinoma infiltration by effector T cells"

**This PDF file contains:**

**Fig. S1-S4**

**Table S1-S2**

**Supplementary Figures**

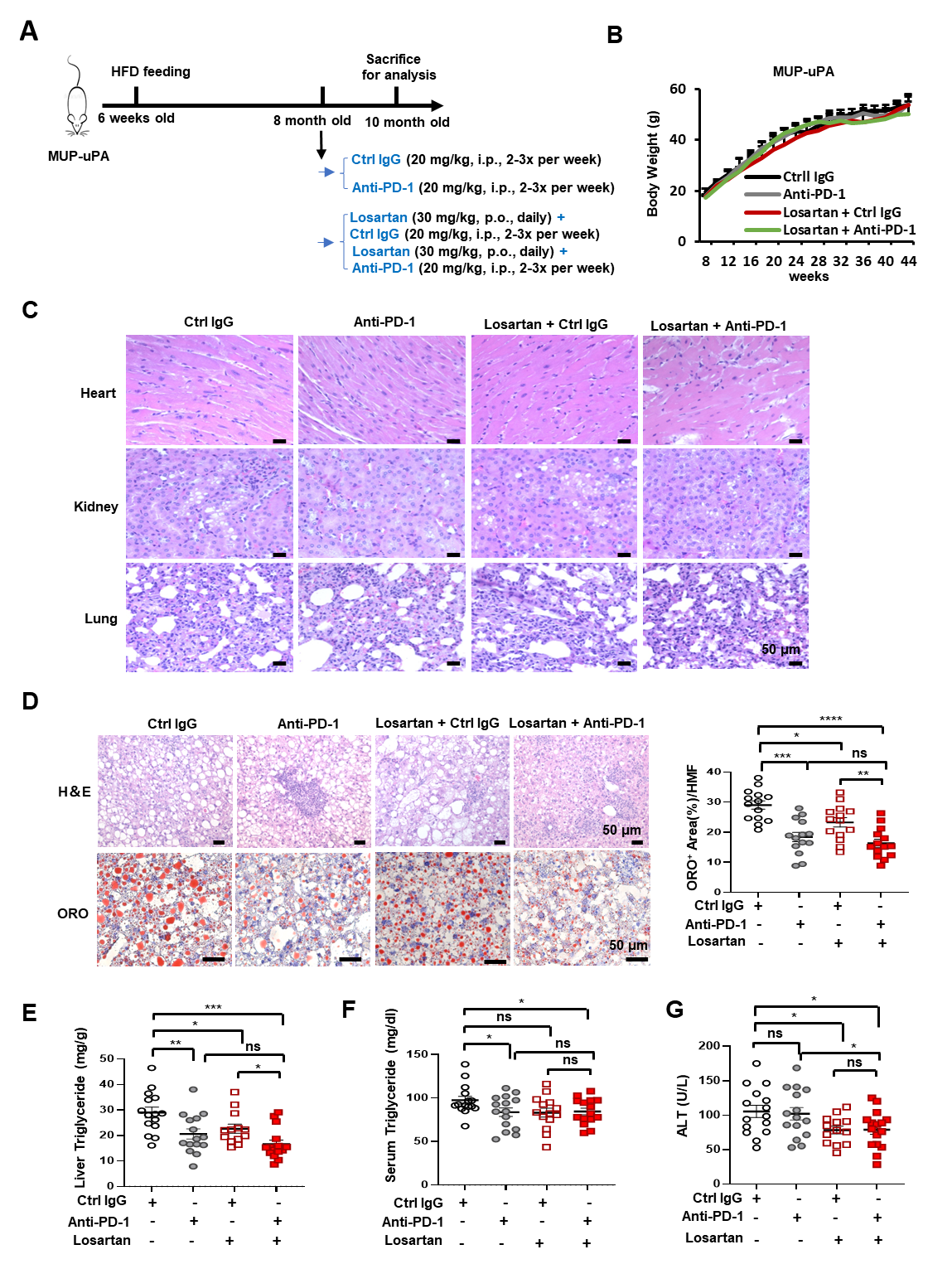

**Fig. S1.** Losartan potentiates anti-PD-1 induced HCC regression.

(*A*) Outline of HFD-induced NASH-HCC in *MUP- uPA* mice and treatment scheme.

(*B*) Body weight gain by HFD-fed *MUP- uPA* mice subjected to the indicated treatments (n=13-15).

(*C*) H&E staining of heart, kidney, and lung tissues of mice from the different treatment groups outlined in S1A (n=13-15). Scale bars, 50 μm.

(*D*) H&E and Oil Red O (ORO) staining of liver sections from the different treatment groups (n=8) (left). Scale bars, 50 μm. ORO staining intensity per high-magnification-field (HMF) determined by Image J (right).

(*E-G*) Liver (E) and serum (F) triglycerides (TG), and serum alanine aminotransferase (ALT) (G) in mice belonging to the different treatment groups.

Data are presented as mean ± SEM. *P < 0.05, **P < 0.01, ***P < 0.001, ****P < 0.0001 (Unpaired two-tailed t test and Mann-Whitney test).

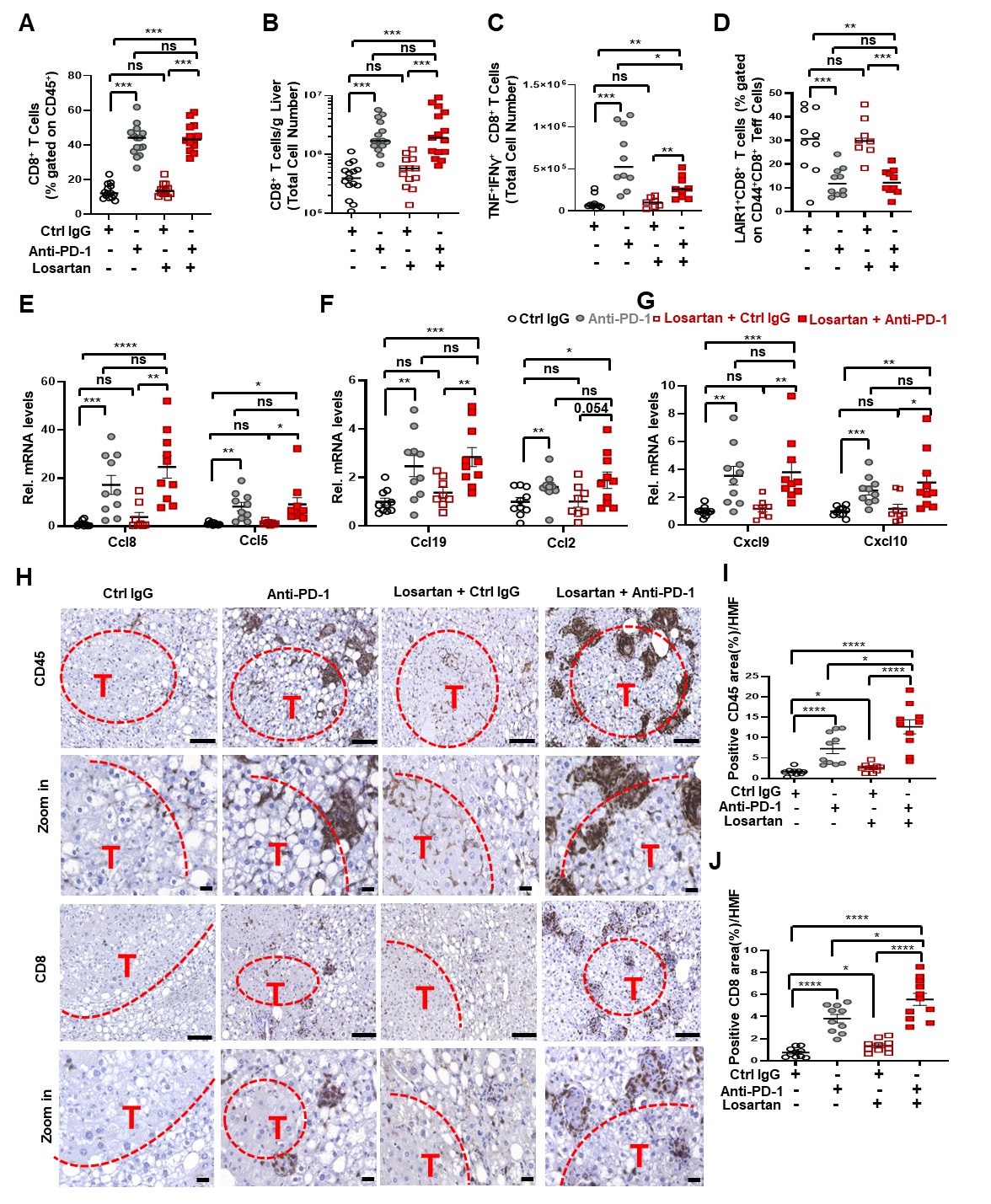

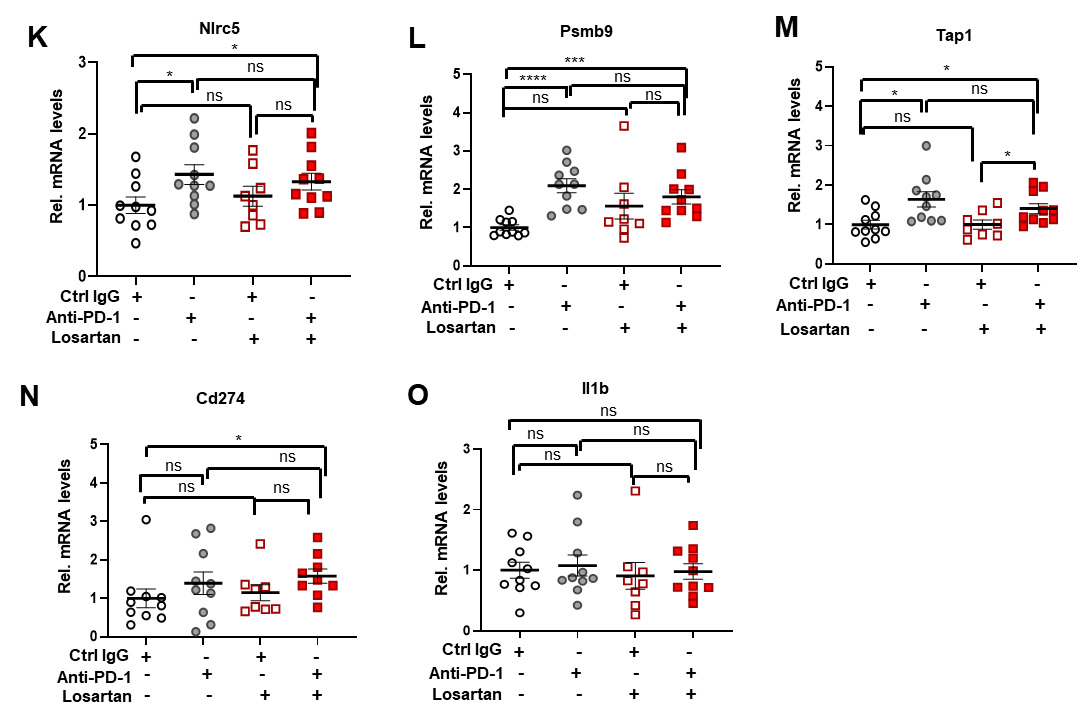

**Fig. S2.** Losartan stimulates intratumoral infiltration of anti-PD-1 reinvigorated Teff cells.

(*A-D*) Liver CD8^+^ T cells as percentage of liver CD45^+^ cells (A), CD8^+^ T cell number/g liver (B), liver TNF^+^IFNγ^+^ CD8^+^ T cell number (C), and LAIR1^+^CD8^+^ cells as percentage of total liver CD44^+^CD8^+^ Teff cells (D) determined by FC analysis of liver cell suspensions from the different treatment groups (n=13-15).

(*E-G*) Relative mRNA amounts of Ccl8, Ccl5 (E), Ccl19, Ccl2 (F), Cxcl9, and Cxcl10 (G) determined by Q-RT-PCR analysis of liver RNA from the indicated treatment groups.

(*H*) FFPE liver sections from the different treatment groups stained for CD45 and CD8 (n=8). T-tumor, scale bars, 50 μm.

(*I* and *J*) Quantification of CD45 (I), and CD8 (J) positive areas per HMF determined by Image J.

(*K-O*) Q-RT-PCR quantitation of Nlrc5 (K), Psmb9 (L), Tap1 (M), Cd274 (N), and Il1b (O) mRNAs in livers from the indicated treatment groups.

Data are presented as mean ± SEM. *P < 0.05, **P < 0.01, ***P < 0.001, ****P < 0.0001 (Unpaired two-tailed t test and Mann-Whitney test).

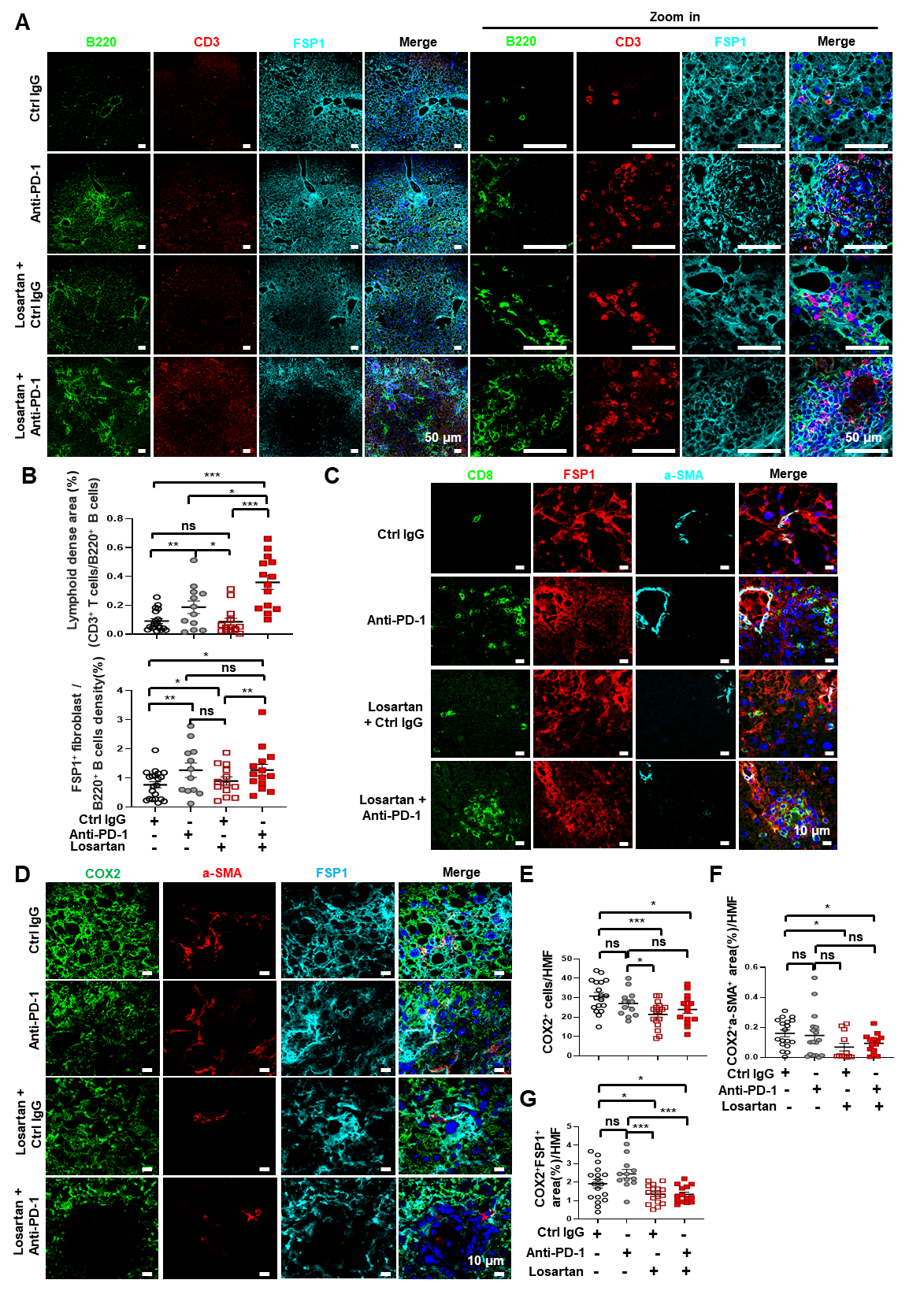

**Fig. S3.** Losartan decreases liver fibrosis induced by anti-PD-1 treatment.

(*A*) Frozen liver sections from the indicated treatment groups were stained for B220, CD3, and FSP1 (n=6-8). Scale bars, 50 μm.

(*B*) Lymphoid dense areas (top) and FSP1^+^ fibroblast/B220^+^ B cell density (bottom) per HMF determined by Image J.

(*C*) Frozen liver sections from the indicated treatment groups were stained for CD8, FSP1, and αSMA and examined by fluorescence microscopy (n=6-8). Scale bars, 10 μm. Experiments were repeated at least three times.

(*D*) Frozen liver sections from the indicated treatment groups were stained for COX2, αSMA, and FSP1 (n=6-8). Scale bars, 10 μm.

(*E-G*) Quantification of areas occupied by COX2^+^ (E), COX2^+^αSMA^+^ (F), and COX2^+^FSP1^+^ (G) per HMF from the images shown in Fig. S3D.

Data are presented as mean ± SEM. *P < 0.05, **P < 0.01, ***P < 0.001 (Unpaired two-tailed t test and Mann-Whitney test).

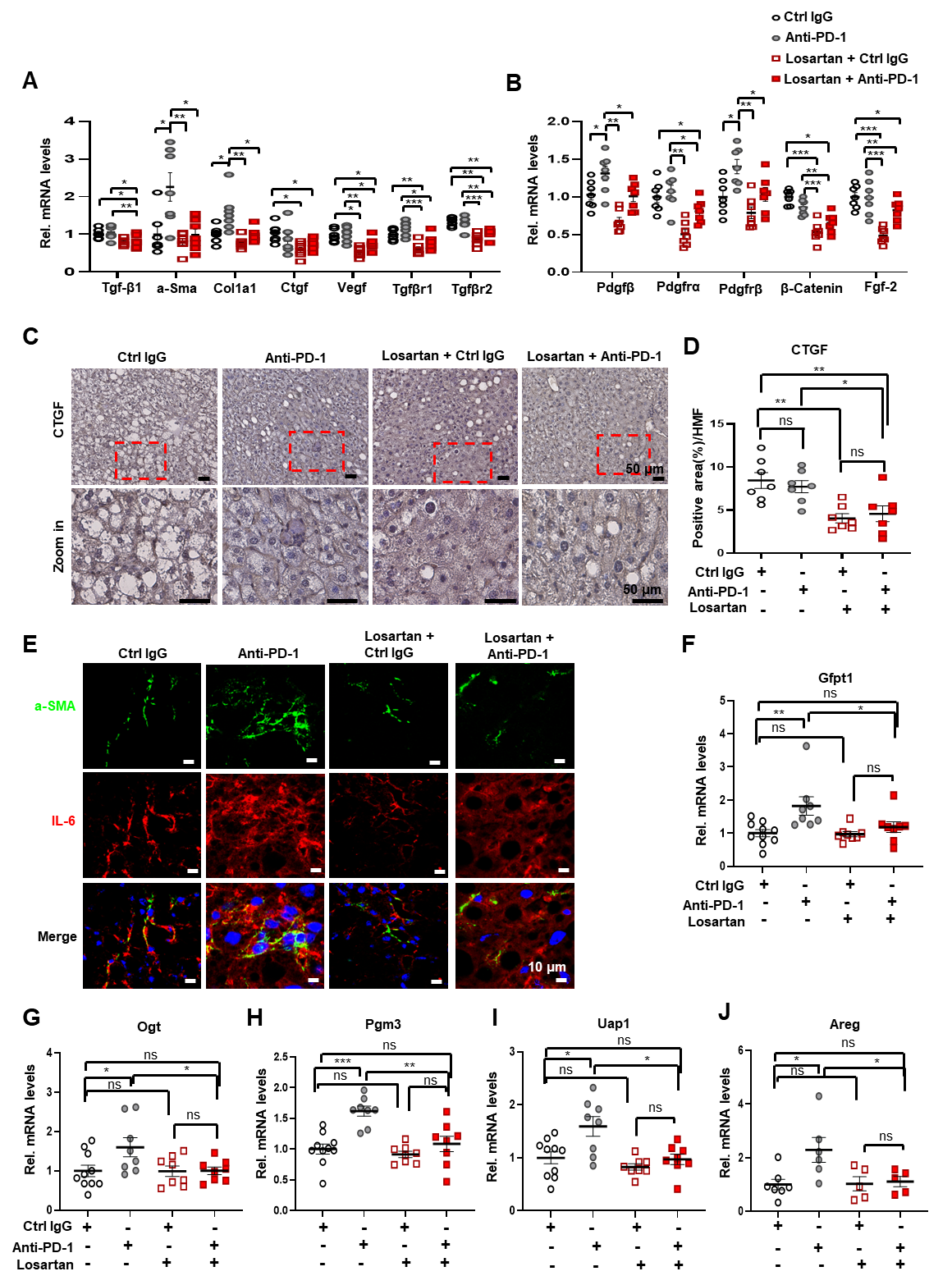

**Fig. S4.** Losartan promotes stroma remodeling, and represses the hexosamine biosynthetic pathway.

(*A* and *B*) Q-RT-PCR quantitation of mRNAs related to TGF-β (A) and PDGF (B) signaling in livers from the indicated treatment groups (n=7).

(*C* and *D*) FFPE liver sections from the different treatment groups were stained for CTGF (C) (n=8), whose staining intensity (D) per HMF was determined by Image J. Scale bars, 50 µm.

(*E*) Frozen liver sections from the different treatment groups were stained for αSMA and IL-6 and examined by fluorescence microscopy (n=6-8). Scale bars, 10 µm.

(*F-J*) Q-RT-PCR quantitation of mRNAs coding for the hexosamine biosynthetic pathway (F-I) and Areg (J) in livers from the indicated treatment groups (n=8-10).

Data are presented as mean ± SEM. *P < 0.05, **P < 0.01, ***P < 0.001 (Unpaired two-tailed t test and Mann-Whitney test).

**Table S1.** Antibodies used in this study.

| Antibodies to: | Catalog number | Source |  |
| --- | --- | --- | --- |
| p-SMAD2(S465/467)/  SMAD3(S423/425) | #8828 | Cell Signaling | Rabbit |
| CD45 | #70257 | Cell Signaling | Rabbit |
| FSP1 | ab197896 | Abcam | Rabbit |
| CD3 | IS503 | Dako | Rabbit |
| COL1A1 | 72026 | CST | Rabbit |
| COL1A1 | sc8784 | Santa Cruze | Goat |
| α-SMA | #19245 | CST | Rabbit |
| TGFβ | SC130348 | Santa Cruze | Mouse |
| P-ERK1/2 | 9101 | Cell Signaling | Rabbit |
| P-SAMD2 | 18338 | CST | Rabbit |
| P-SMAD3 | 9520 | CST | Rabbit |
| SMAD2/3 | 8685 | CST | Rabbit |
| ERK1/2 | 9102 | CST | Rabbit |
| IL-6 | sc-28343 | Santa Cruze | Mouse |
| B220 | 13-0452-86 | eBioscience | Rat |
| COX2 | PA1-9032 | Thermo Fisher | Goat |
| P21 | ab188224 | Abcam | Rabbit |
| P16 | sc-1661 | Santa Cruz Biotechnology | Mouse |
| CD8 | #98941 | Cell Signaling | Rabbit |
| CD8 | 14-0081-85 | eBioscience | Rat |
| IFNγ | 50-7311-82 | eBioscience | Rat |
| IL-17 | 25-7177-80 | eBioscience | Rat |
| CD4 | 48-0042-80 | eBioscience | Rat |
| CD19 | 50-0193-82 | eBioscience | Rat |
| IgA | 11-4204-83 | eBioscience | Rat |
| CD8 | 45-0081-82 | eBioscience | Rat |
| TNF | 11-7321-41 | eBioscience | Rat |
| B220 | 48-0452-82 | eBioscience | Rat |
| TIM3 | 134006 | Biolegend | Rat |
| CD44 | 48-0441-82 | eBioscience | Rat |
| CD45 | 103137 | Biolegend | Rat |
| PD-L1 | 25-5982-82 | eBioscience | Rat |
| CD138 | 142504 | Biolegend | Rat |
| LAIR1 | 12-3051-82 | eBioscience | Armenian hamster |

**Table S2.** Quantitative PCR primers used in this study.

| **mRNA** | **Forward Primer (5’-3’)** | **Reverse Primer (5’-3’)** |
| --- | --- | --- |
| Psmb9 | GAAGAAGTCCACACCGGGAC | GAGGGGAGAGCTTGTCGAAC |
| Cd274 | GCTCCAAAGGACTTGTACGTG | TGATCTGAAGGGCAGCATTTC |
| Nlrc5 | GACGCTGGGGTTAACAGGAA | CAGCTCCACAAGACTCAGCA |
| Tap1 | CCCAGCAGGTTCCATCACAT | GAAAAAGCAGGGGCAGGTTG |
| IL1b | GCCTCGTGCTGTCGGACC | TGTCGTTGCTTGGTTCTCCTTG |
| Ccl8 | TCTACGCAGTGCTTCTTTGCC | AAGGGGGATCTTCAGCTTTAGTA |
| Ccl19 | GGGGTGCTAATGATGCGGAA | CCTTAGTGTGGTGAACACAACA |
| Ccl5 | GCTGCTTTGCCTACCTCTCC | TCGAGTGACAAACACGACTGC |
| Ccl2 | TTAAAAACCTGGATCGGAACCAA | GCATTAGCTTCAGATTTACGGGT |
| Cxcl9 | TGCCATGAAGTCCGCTGTTC | CTAGGGTTCCTCGAACTCCAC |
| Cxcl10 | CCAAGTGCTGCCGTCATTTT | TTCATCGTGGCAATGATCTCAAC |
| α-Sma | CTGACAGAGGCACCACTGAA | GAAGGAATAGCCACGCTCAG |
| Tgf-β1 | TTGCTTCAGCTCCACAGAGA | TGGTTGTAGAGGGCAAGGAC |
| Col1a1 | GCTCCTCTTAGGGGCCACT | CCACGTCTCACCATTGGGG |
| Ctgf | CAAAGCAGCTGCAAATACCA | GTCTGGGCCAAATGTGTCTT |
| Tgf-βrI | GGTCTTGCCCATCTTCACAT | CAGGGGCCATGTACCTTTTA |
| Tgf-βrII | GCAAGTTTTGCGATGTGAGA | GGCATCTTCCAGAGTGAAGC |
| Vegf | CAAGATCCGCAGACGTGTAA | TTAATCGGTCTTTCCGGTGA |
| Pdgfrα | TGGCATGATGGTCGATTCTA | CGCTGAGGTGGTAGAAGGAG |
| Pdgfrβ | TCAACGACTCACCAGTGCTC | TTCACAGGCAGGTAGGTGCT |
| Pdgfβ | TCCAGATCTCTCGGAACCTC | GGCTTCTTTCGCACAATCTC |
| β-Catenin | CTCTTCAGGACAGAGCCAATG | ATGCTCCATCATAGGGTCCA |
| Fgf-2 | CCTTGCTATGAAGGAAGATGG | TCCGTGACCGGTAAGTATTG |
| Ogt | GACGCAACCAAACTTTGCAGT | TCAAGGGTGACAGCCTTTTCA |
| Pgm3 | AGCAGTGGGATGCTATTTATGTC | TGTCTGCGCTTTCTTGTGAGT |
| Uap1 | ACTCCCAGGGGCACTTCATTA | GGCCTCTGACTTTCCATTTGT |
| Areg | TCATGGCGAATGCAGATACA | GCTACTACTGCAATCTTGGA |
| Gfpt1 | GAAGCCAACGCCTGCAAAATC | CCAACGGGTATGAGCTATTCC |
